# A Robust and Scalable Workflow for the Production of Circular Single-Stranded DNA for Genome Engineering Applications

**DOI:** 10.64898/2026.08.14.743880

**Authors:** Susi Mathews, Mahima Kapoor, Akschya Sivacoumar, Raj Acharya, Souvik Maiti, Debojyoti Chakraborty

## Abstract

Circular single-stranded DNA (cssDNA) is a versatile biomolecule with applications spanning genome editing, DNA nanotechnology, synthetic biology, molecular diagnostics, and aptamer development. Compared with linear single-stranded DNA, cssDNA offers enhanced structural stability, resistance to in-cellulo degradation by exonucleases and enables the generation of long, sequence-defined DNA molecules that are difficult to obtain through conventional chemical synthesis methods. Despite its growing utility, widespread adoption of cssDNA has been limited by the lack of accessible, scalable, and cost-effective production methods, with many existing workflows relying on specialised reagents, extensive optimisation, or commercially synthesised DNA.

Here, we present a streamlined, end-to-end protocol for the laboratory-scale production of high-purity cssDNA using an M13 phagemid-based system and standard molecular biology laboratory infrastructure. The workflow encompasses bacterial culture, phage amplification, nuclease treatment, phage precipitation, anion-exchange purification, and quality control, with practical optimisations to improve yield, reproducibility, and scalability. Using this approach, yields range from 120–195 µg of purified cssDNA from 300 mL of culture supernatant. The protocol provides detailed guidance on critical process parameters, troubleshooting, and quality assessment, enabling reliable production of cssDNA suitable for a wide range of downstream molecular biology and genome engineering applications.

## Introduction

Filamentous phage (e.g., M13/f1/fd) systems enable low-cost biological production of cssDNA by coupling an f1/M13 origin of replication and packaging signal to secretion of ssDNA-containing bacteriophages from *E. coli* into culture supernatants [1,2]. We have developed a streamlined, end-to-end workflow for cssDNA production, adapted in part from previously published phage-derived ssDNA purification protocols and methodologies. Key modifications were introduced to improve yield, reproducibility, and scalability, making high-quality cssDNA production more accessible to standard molecular biology laboratories.

### Principle of phage-based cssDNA production

Phage-based production of circular single stranded DNA (cssDNA) relies on the non-lytic replication and secretion cycle of filamentous bacteriophages of the Ff class, including M13, f1, and fd. Unlike lytic phages, filamentous phages establish a chronic infection in Escherichia coli, during which phage genomes are continuously replicated and secreted from the cell without host lysis. This life cycle is in particular characterised by the generation and packaging of covalently closed circular ssDNA, which represents the native genomic form of filamentous phages [3,4,5].

The essential cis-acting element required for cssDNA production is the origin of replication of filamentous phages i.e f1 ori. Constructs containing this origin, such as phagemids or isogenic miniphages, are maintained in the cells as double-stranded circular replicative forms. Upon expression of phage replication proteins supplied in trans, the gene II protein (pII) specifically recognizes the f1 origin and introduces a site-specific nick in the positive (+) strand. This nick creates a free 3′-hydroxyl group that primes host DNA polymerase to initiate rolling-circle replication, resulting in release of the parental (+) strand as single-stranded DNA. A second pII-mediated cleavage-ligation event releases the covalently closed circular ssDNA molecule [4,6].

The newly synthesised ssDNA is rapidly coated by the gene V protein (pV), a high-affinity single-stranded DNA-binding protein that shields ssDNA from nucleolytic degradation and prevents its re-annealing to complementary strands. Binding by pV also commits the ssDNA to the phage assembly pathway, re-directing it away from being converted back to double-stranded DNA [5,7]. The ssDNA-pV complex is then taken to membrane-embedded assembly sites, where ssDNA extrusion and virion assembly occur simultaneously. During this process, pV is displaced by the major coat protein pVIII, yielding filamentous virions that each contain a single molecule of circular ssDNA.

Current cssDNA production platforms unlink the circular ssDNA to be packaged from the phage replication and assembly machinery by supplying phage proteins on helper plasmids lacking f1 origin. This design ensures that only the f1-containing construct is replicated as cssDNA and packaged, thus facilitating scalable production of pure cssDNA for biotechnological and therapeutic applications [8,9,10].

**Figure 1:**
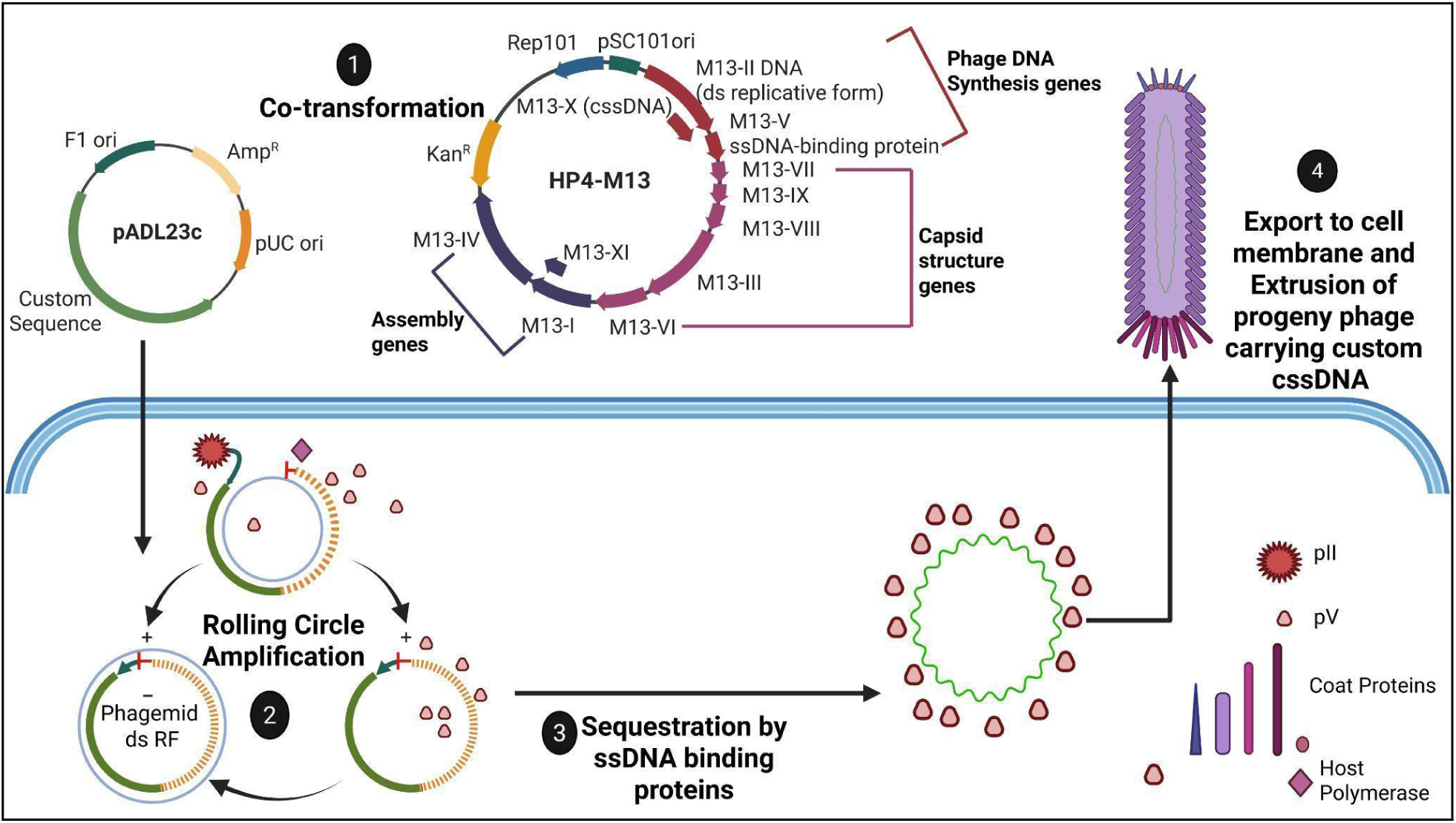
Principle of Phage-based circular single-stranded DNA Production. *(Generated via Biorender)* *1) Co-transformation of F1 pilus+ bacterial host by a) a phagemid having an M13 replication ori along with an insert of interest and b) a helper-phage plasmid encoding all necessary phage proteins. 2) Expression of phage proteins in trans helps carryout out rolling circle amplification of the double-stranded circular replicative forms (RF) phagemid leading to formation of single-stranded DNA (plus strand) 3) Sequestration of cssDNA by Phage ssDNA binding proteins (pV) re-routes the replication cycle towards assembly-extrusion. 4) Extrusion of Progeny phage particles containing custom cssDNA*.

### Potential Applications of cssDNA

Circular single-stranded DNA (cssDNA) is a practical nucleic acid format that combines single-strandedness with covalently closed circular topology (i.e., no free ends). This makes it an excellent template for many enzymatic reactions and provides enhanced resistance to end-directed nucleases. These characteristics have allowed cssDNA to serve as an enabling reagent in the field of molecular biology, supporting applications that rely on precise sequence-directed interactions, including mutagenesis, sequencing, genome engineering and nanotechnology.

Filamentous phage M13 and other related phagemid systems have played an important role in early recombinant DNA cloning and sequencing technology. In these systems, the DNA insert was first introduced into the double-stranded replicative form (dsRF) of the vector which led to the infected bacteria producing phage particles containing the recombinant strand as circular single-stranded DNA [11]. This cssDNA could be used directly as a primer extension template, thereby avoiding the additional strand separation/denaturation steps required for double-stranded DNA. Furthermore, the placement of conserved primer binding sites next to the cloning region allowed the same vector-specific primer to be used for many different inserts [12]. M13-derived cssDNA was subsequently adopted for shotgun sequencing and contributed to the sequencing of human mitochondrial DNA fragments and the complete cauliflower mosaic virus genome [13,14]. Use of oppositely oriented paired M13 vectors also further allowed either strand of a cloned fragment to be recovered and sequenced [15].

cssDNA has also been commonly employed for oligonucleotide-directed mutagenesis using a uracil-containing circular ssDNA template typically produced from M13/phagemid systems. In the original Kunkel protocol, the mutagenic primer was annealed to cssDNA and a complementary strand was synthesised. As the biological selection and repair biases favoured this newly synthesised, covalently closed, mutated, circular DNA, the resultant product carrying intended change was generated. This makes cssDNA an integral substrate for mutagenesis-based work [16,17].

In 2006, Rothemund demonstrated that, DNA origami involved folding of a long ssDNA scaffold (typically the 7kb M13 bacteriophage circular genome) into definite nanoscale shapes via a collection of short, single-stranded “staple” oligonucleotides [18]. The requirement for customised sequence- and size-specific scaffolds that extend origami design space beyond the capabilities of the M13 genome is a primary motivation for development of cssDNA production methods. Bioproduction of kilobase cssDNA in a miniphage format has been shown and applied to nanotechnology applications, including scaffolded DNA origami and sequence-encoded information [8,9].

cssDNA combines the repair-compatible properties of long ssDNA with an end-free topology that can improve donor stability and permit the manufacture of templates beyond the practical size range of synthetic oligonucleotides [9,19,20]. Owing to its circular nature, it could potentially circumvent free-end dependent concatemerisation and rearrangement issues widely linked to unmodified linear dsDNA donors and plasmid dsDNA donors respectively. Reports have also suggested that it is associated with reduced cGAS-mediated innate-immune toxicity in comparison to other forms [9,21].

In the first published experimental study that focused on cssDNA as a Homology Directed Repair (HDR) template, Iyer et al. showed that cssDNA works as an effective HDR template when used with both CRISPR-Cas9 and Cas12a, yielding integration frequencies higher than linear ssDNA donors in their comparative data [19]. Building on the same core idea, Xie et al. introduced the concept of non-viral genome cssDNA donor system called GATALYST (Genome engineering CATALYST), that shows cssDNA as a superior HDR donor compared to dsDNA and lssDNA, highlighting its enhanced integration efficiency, improved safety and the capability to deliver large transgene cargos. GATALYST demonstrated scalable production of cssDNA donors up to ∼20 kb, knock-in efficiencies up to 70% in iPSCs, and improved precise integration across multiple primary immune cell types i.e. primary T, NK and B cells, as well as CD34+ HSPCs; across multiple genomic loci through various nuclease editor platforms [12]. The enGager/TESOGENASE system devised recently has shown to increase cssDNA knock-in by conjugating these donors to nuclear-targeted Cas9 through ssDNA-binding motifs achieving up to ∼6-fold greater integration than those without enGAGER and enabling CAR knock-in rate of ∼33% in primary human T cells [16]. More recently, Letort et al. have extended cssDNA applicability to TALEN editing in HSPCs and demonstrated 3-5× higher knock-in frequency than lssDNA with efficiencies greater than 40%. They also reported superior engraftment and maintenance of edits compared to AAV-edited controls in a murine model [20]. All of these studies have led to cssDNA being considered a promising donor format for gene editing workflows.

In biosensing and diagnostics, rolling circle amplification (RCA) employs cssDNA templates to generate long concatemeric products for highly sensitive target detection and signal amplification. Standardised, high-quality cssDNA can therefore improve reproducibility and reduce background in diagnostic platforms [22,23]. Circularization also enhances aptamer performance by removing free ends that are vulnerable to nuclease degradation in biological fluids [24,25]. Circular DNA aptamers have been evolved directly in serum while retaining strong target binding and improved biostability [19]. In addition, compared to their non-circular bivalent counterpart, circular bivalent aptamers exhibited improved thermal stability, resistance to degradation by nucleases, and overall greater accumulation/retention in tumours as determined by in vivo fluorescence imaging [25].

Beyond acting as a scaffold or repair template, biologically produced cssDNA could also be potentially used as a sequence-defined substrate for information encoding (digital data stored in DNA sequence) coupled to biological amplification, illustrating its possible value as both a physical biomaterial and an information carrier [8].

**Figure 2:**
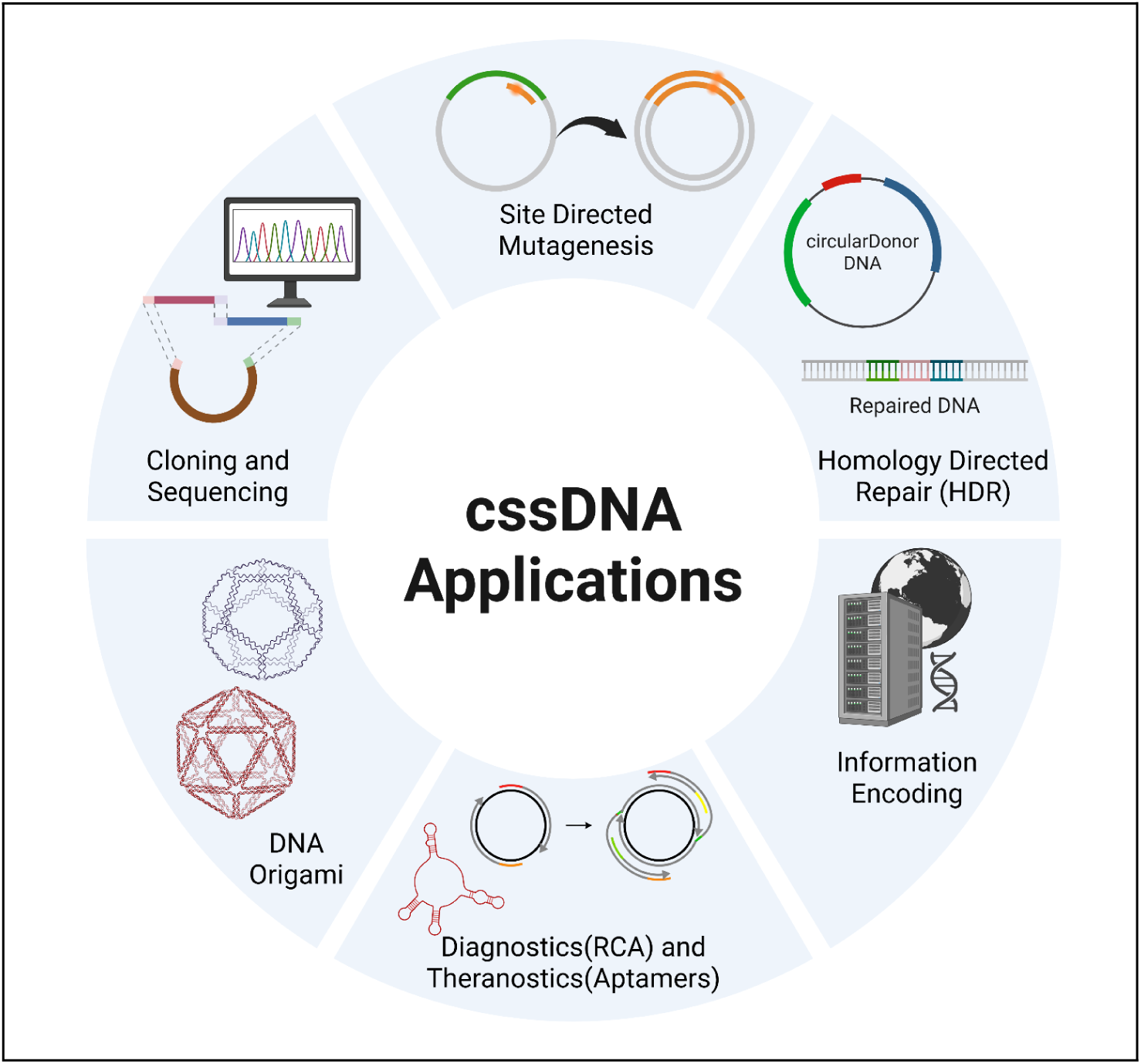
Applications of cssDNA in various fields. *(Generated via Biorender)*

### Drawbacks of existing protocols

Despite the widespread use of filamentous phage–based systems for cssDNA production, several practical challenges limit their robustness and ease of adoption in routine laboratory workflows. A recurring difficulty is the requirement for co-transformation or co-maintenance of a low-copy helper plasmid (e.g., M13-derived constructs) alongside a higher-copy phagemid. Because helper plasmids are often maintained at substantially lower copy numbers, co-transformation efficiency can be variable and frequently necessitates empirical optimisation of DNA input ratios, with higher relative amounts of helper plasmid required to achieve stable dual maintenance. This variability can result in inconsistent phage production and increased experimental repetition, particularly for users unfamiliar with phagemid systems [8,26,27].

In addition, many existing protocols provide limited guidance on culture growth dynamics and harvest timing, which can complicate reproducibility and workflow planning. Inoculation of secondary cultures from overnight starters frequently results in rapid increases in optical density, causing cultures to reach optimal harvest points at unpredictable or inconvenient time intervals. This lack of temporal control can hinder the introduction of logical pause points in the workflow and may lead to either premature harvesting, reducing phage yield, or delayed harvesting, which is often associated with increased host cell stress and release of contaminating nucleic acids [1,2,28].

Finally, several existing methods rely on small culture volumes or tightly time-coupled batch workflows that are poorly aligned with routine laboratory schedules. Continuous monitoring is often required to capture optimal growth phases, limiting scalability and increasing operator burden. Collectively, these factors contribute to batch-to-batch variability, elevated hands-on time, and reduced accessibility for laboratories seeking to implement cssDNA production at preparative scale [8,29].

### Brief Overview of the Current Protocol

In this streamlined protocol, we utilise a phage-based phagemid system to generate circular single-stranded DNA (cssDNA) containing custom sequence inserts, leveraging the ability of filamentous M13 bacteriophage f1 origins to support rolling circle replication and export of the M13 phage containing ssDNA into the culture medium. Phagemids harboring the desired sequence are co-transformed into an f+ *E. coli* (XL1 Blue) host along with an M13 HP4 helper plasmid obtained from Addgene (Plasmid #120340) that supplies the phage replication and packaging functions in trans, enabling preferential packaging and secretion of the target cssDNA while minimising incorporation of helper sequences into the final product. This strategy has been widely used for the production of ssDNA scaffolds in applications ranging from DNA origami to programmable vectors for gene expression, with optimised systems achieving milligram-scale yields and high sequence purity [30]. After exponential culture growth, phage particles are harvested from the filtered supernatant, treated with DNase I to degrade contaminating double-stranded DNA from lysed host bacterial cells, and purified using Qiagen-tip 500. The resulting cssDNA is assessed for yield and quality by spectrophotometry and agarose gel electrophoresis prior to downstream use. By tuning culture conditions and processing volumes, this approach provides a flexible, scalable route to produce high-quality cssDNA for both structural and functional molecular biology applications [20,30].

**Table 1.**
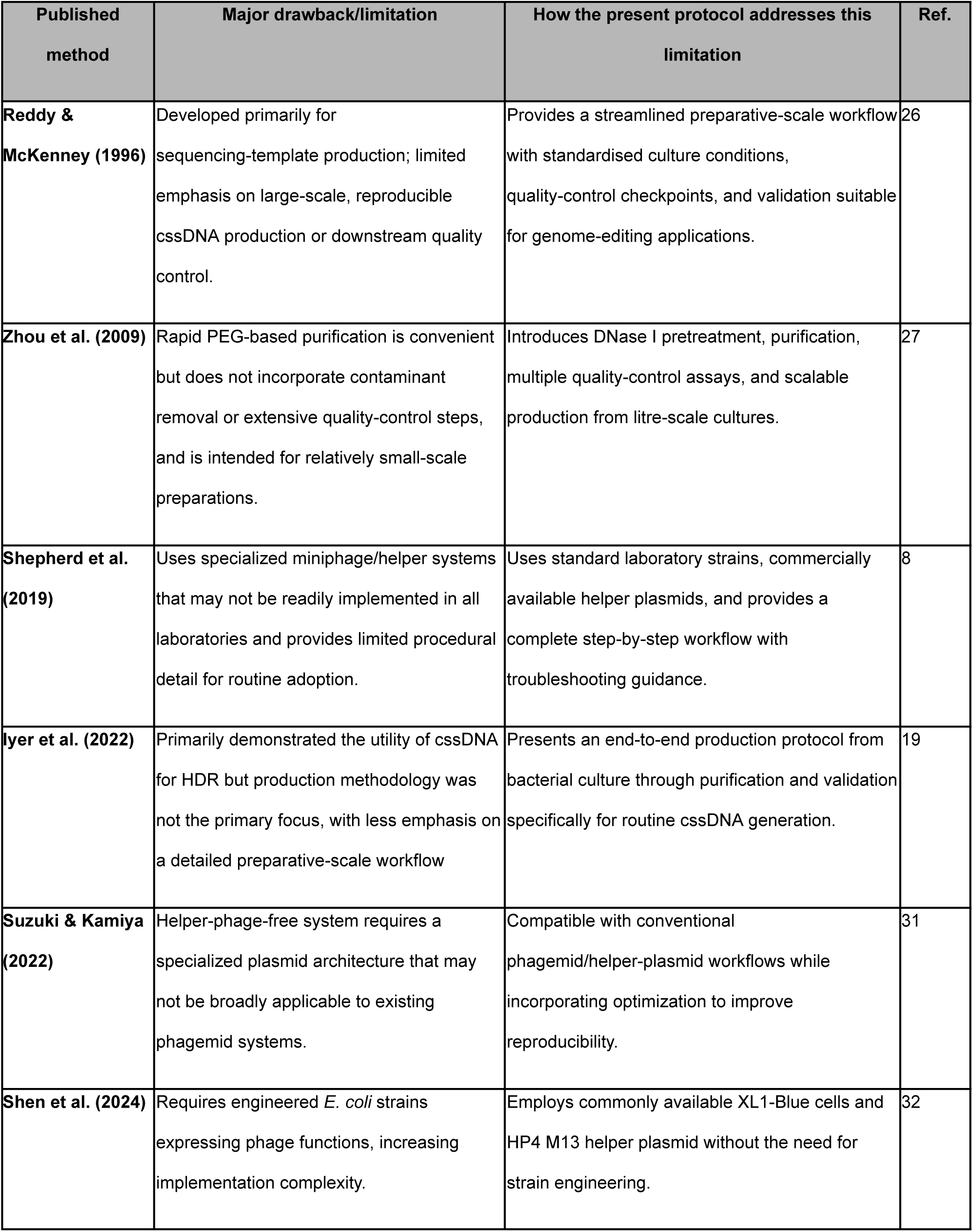
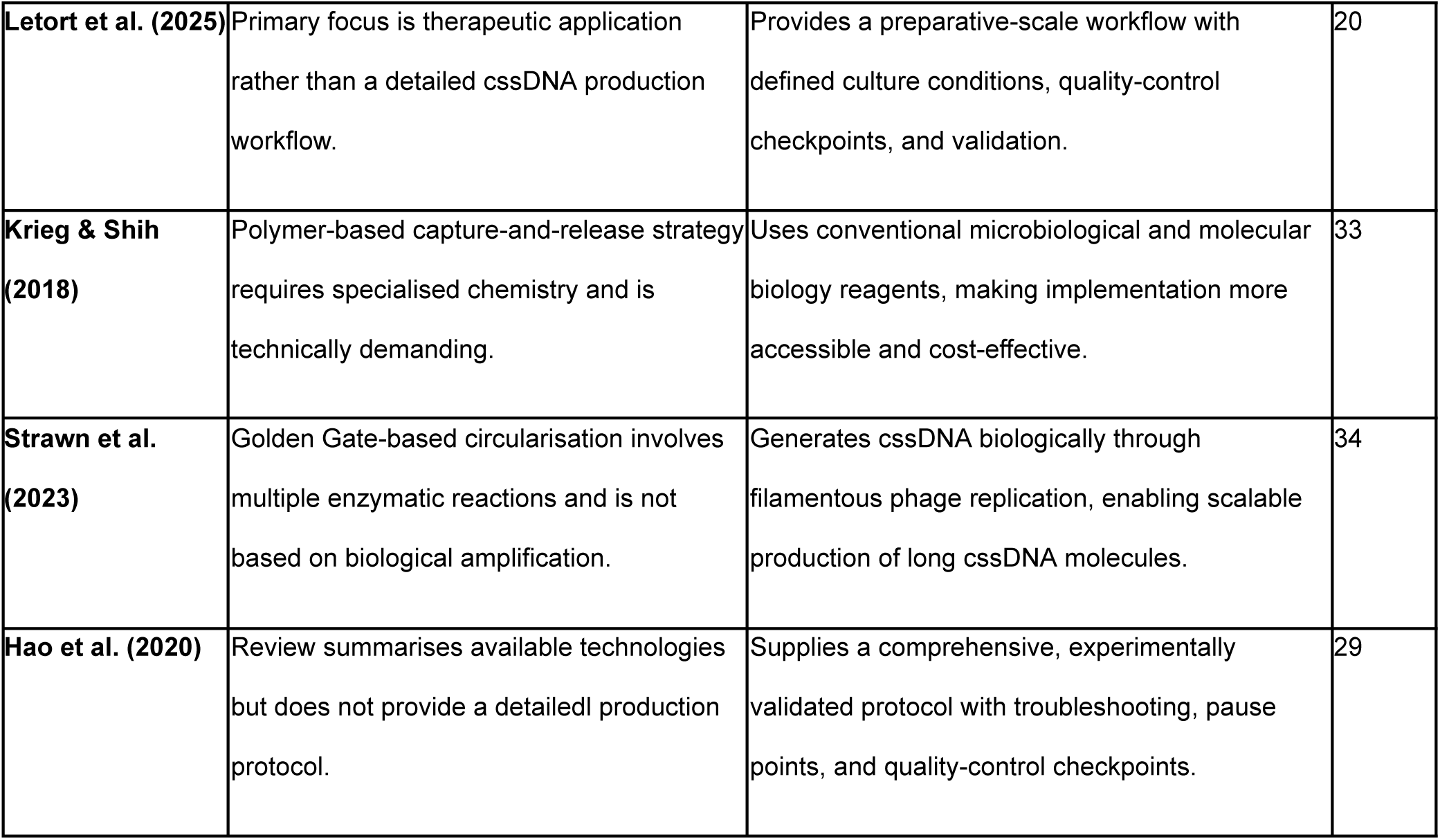
Comparison of the present protocol with previously published cssDNA production protocols.

### Need for Scalable cssDNA Production

Long single-stranded DNA is useful in the field of DNA nanotechnology and CRISPR-based genome editing; however, its production at scale raises challenges because chemical synthesis is length-limited and enzymatic routes often require complex processing or generate mixed products [19]. Biological production of cssDNA using filamentous phage has emerged as a scalable alternative for producing sequence-defined ssDNA at milligram scales under optimised conditions [19]. In filamentous phage biology, plus-strand synthesis initiates and terminates at the f1 origin, which contains overlapping functional domains controlling initiation and termination of viral strand synthesis [1,8]. This origin also functions as a signal enabling preferential packaging of ssDNA derived from a phagemid bearing an f1/M13 origin when helper functions are supplied in trans.

Classic helper-phage approaches (e.g., M13KO7) are widely used to generate secreted phagemid ssDNA from *E. coli* cultures, but can co-produce helper-genome contaminants that require purification or process engineering to minimize. Recent systems reduce contamination by replacing helper phage with engineered helper plasmids, helper strains with integrated phage genes, or helper-free controllable systems, improving purity and simplifying processing [2,8,31,32].

Because cssDNA donors can outperform linear ssDNA donors in specific genome editing contexts, there is a growing need for reliable cssDNA production workflows that goes beyond their traditional use in nanotechnology and M13-based sequencing [8,19,30]. Here, we present a standardised, end-to-end and user friendly protocol that addresses the increasing demand for high purity, high yield cssDNA across various applications.

## Results

### Development of an end-to-end phage-based cssDNA production workflow

We established an end-to-end workflow for the production and isolation of cssDNA using a phage-based system (Fig. 3). The workflow involved co-transformation of an M13 helper plasmid with a phagemid carrying the gene of interest (GOI) into competent XL1-Blue cells followed by expansion of the co-transformed culture. Following culture harvesting, the phage-containing supernatant was clarified and filtered prior to nuclease treatment. DNase I treatment was then used to degrade residual free nucleic acids released from lysed bacterial cells while leaving the phage-encapsidated ssDNA protected from digestion. The clarified and nuclease-treated supernatant was then subjected to PEG/NaCl precipitation to concentrate the phage particles. Following phage lysis, cssDNA was extracted using two alternative purification approaches: anion-exchange chromatography using a QIAGEN-tip column and an organic extraction method involving Proteinase K and Phenol Chloroform Isoamyl treatment. The inclusion of two purification strategies provides flexibility in downstream processing, allowing this cssDNA production protocol to be adapted according to the availability of purification columns, reagents, and laboratory infrastructure. Together, these steps constituted a streamlined workflow for the preparative-scale production and recovery of cssDNA.

**Figure 3.**
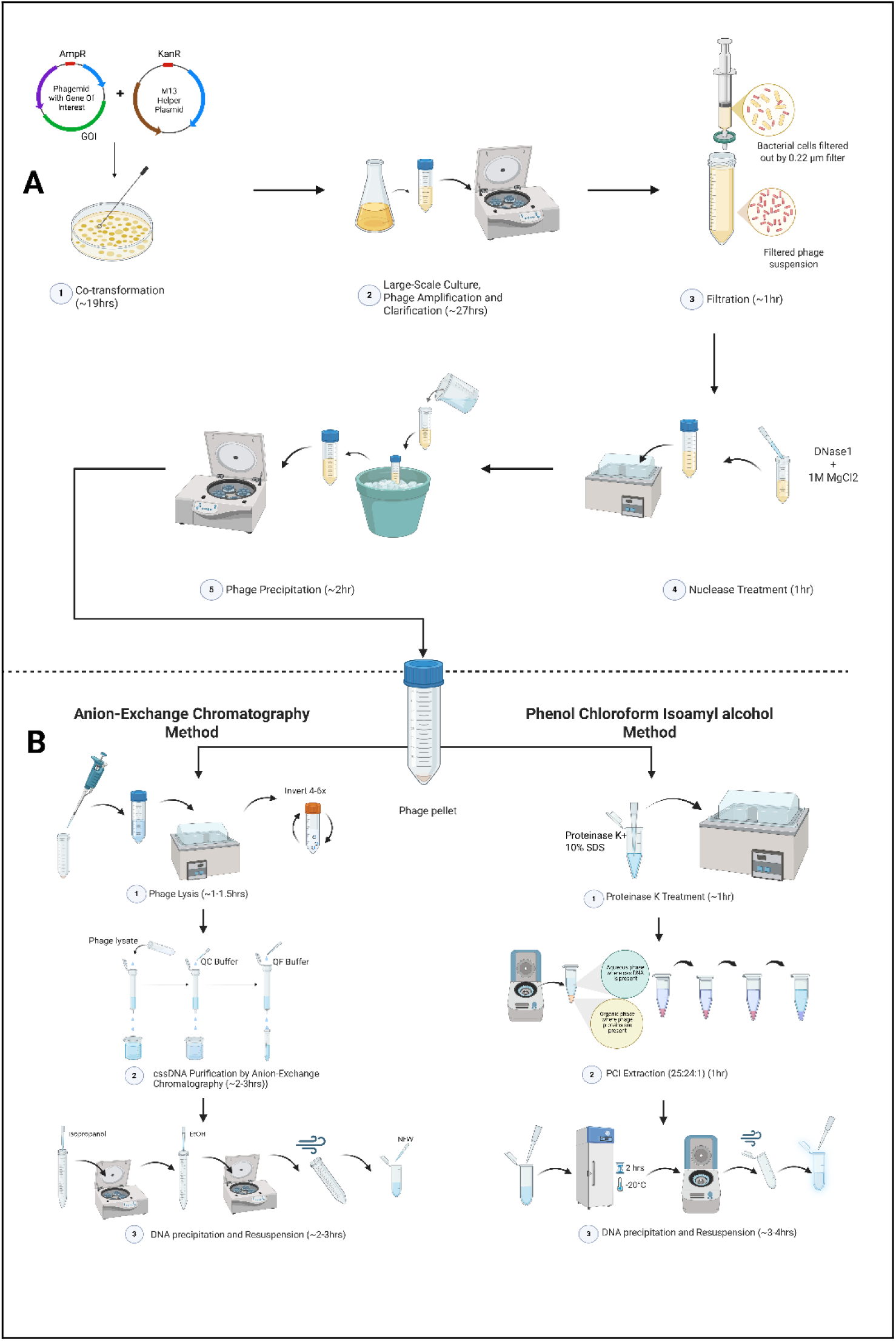
Production of phage particles and downstream processing. **(A)** Workflow for the production of filamentous phage particles in bacterial culture **(B)** Two methods for the isolation and purification of cssDNA from phage particles *(Generated via Biorender)*

Following phage production and culture harvest, we assessed whether supernatant clarification by membrane filtration and nuclease treatment reduced extracellular plasmid DNA present in the culture supernatant. Samples collected before and after filtration and DNase I treatment were analysed by agarose gel electrophoresis. A distinct plasmid-derived band was observed in the unfiltered and untreated supernatants but was substantially reduced or absent following filtration and subsequent DNase I treatment (Fig. 4). These observations supported the inclusion of both filtration and nuclease treatment as critical upstream purification steps for reducing contaminating bacterial DNA prior to phage precipitation and lysis.

**Figure 4.**
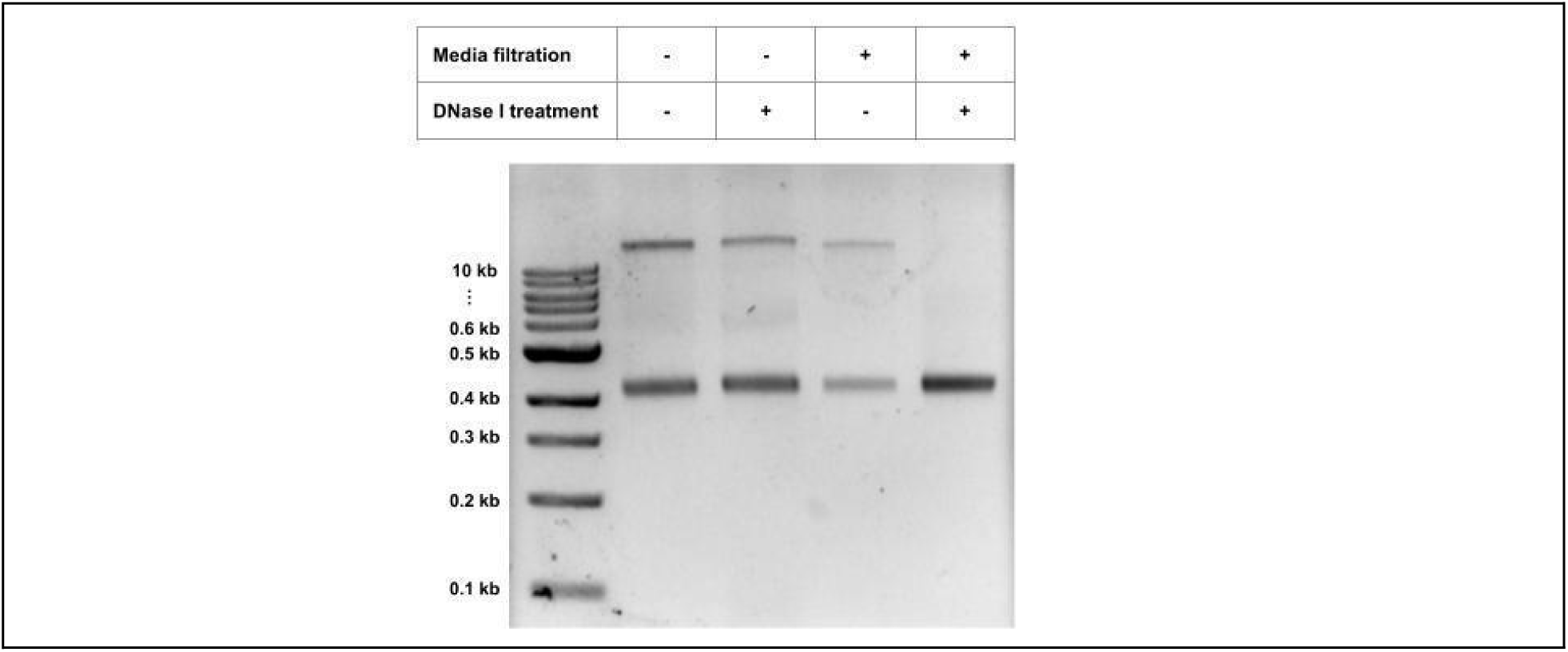
cssDNA isolates with and without filtration and DNaseI treatment. Dual Treatment Reduces Bacterial Plasmid Contamination in Culture Supernatants.

### In-process monitoring of cssDNA production

Prior to large-scale purification, a small aliquot of the bacterial pellet was subjected to small-scale plasmid DNA extraction and analysed by agarose gel electrophoresis to verify the presence of the cssDNA band of the expected size (Fig. 5a). Although the plasmid miniprep was not intended for cssDNA purification, detection of the characteristic cssDNA band served as a rapid in-process quality check, providing confidence that the large-scale culture was producing cssDNA and helping determine whether to proceed with downstream purification. Depending on the construct, bands corresponding to the phagemid, helper plasmid, and cssDNA were observed.

**Figure 5.**
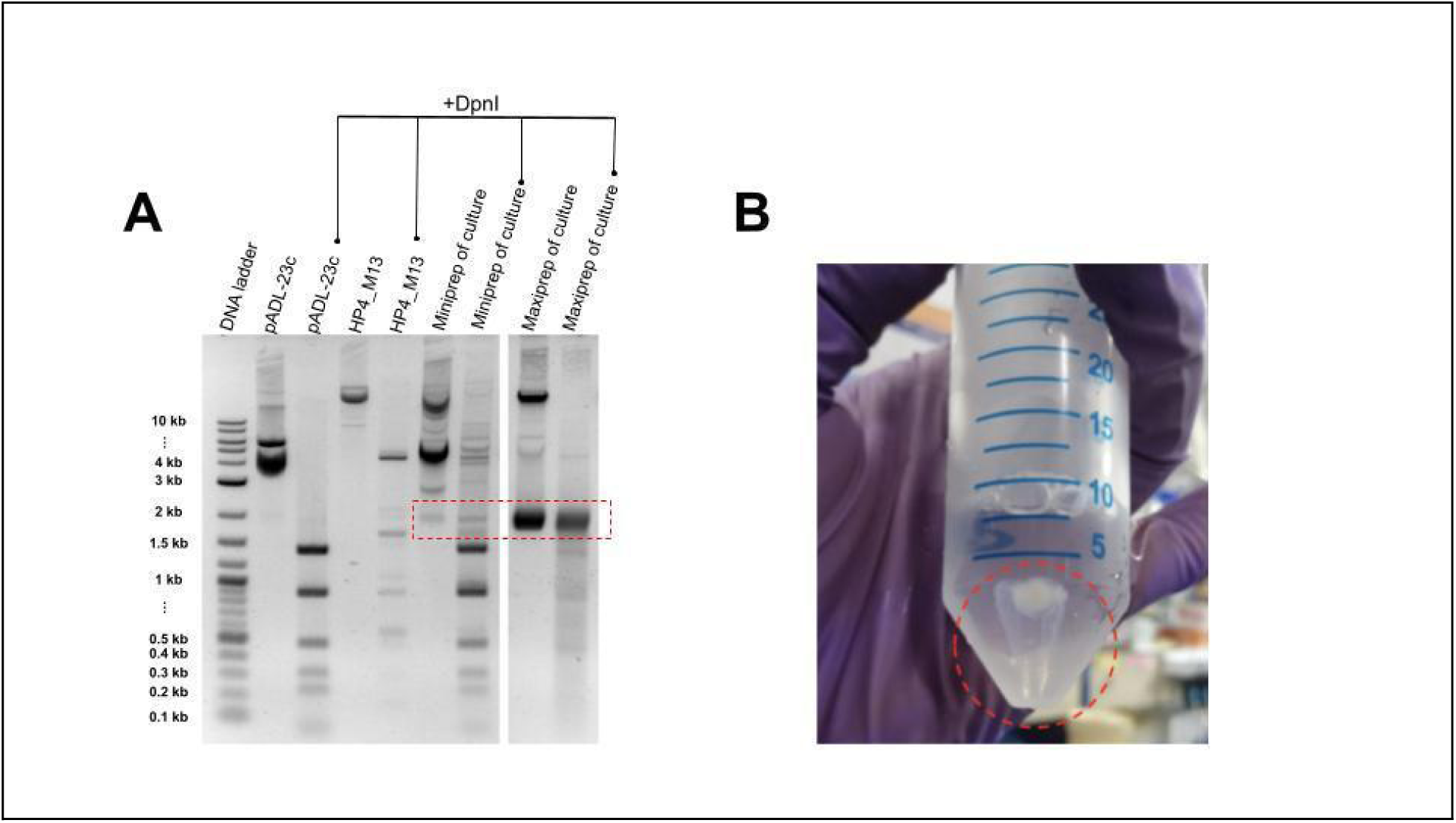
In-process quality checks during phage production. **(A)** Miniprep analysis of the bacterial culture prior to phage isolation can be used to verify cssDNA production. Successful cssDNA synthesis is indicated by the presence of an additional DNA band, distinct from the two co-transformed plasmids. Digestion with DpnI was used to assess the contribution of methylated plasmid-derived DNA to the observed bands. **(B)** Following PEG/NaCl-mediated precipitation and centrifugation of the culture supernatant, successful phage precipitation is indicated by the formation of a visible whitish pellet corresponding to the precipitated phage particles.

### Characterisation of purified cssDNA

Purified cssDNA preparations from four constructs were analysed by agarose gel electrophoresis to assess integrity and size distribution. Distinct bands corresponding to the expected cssDNA products were observed across constructs, with limited evidence of degradation (Fig. 5A).

The circular nature of the isolated DNA can be assessed through enzymatic digestion assays, including resistance to exonucleases that selectively degrade linear single-stranded DNA (e.g., Exonuclease I), while remaining sensitive to treatments that linearise or nick circular DNA. No detectable degradation of the purified isolate was observed, whereas the linear ssDNA control was degraded, supporting the presence of predominantly circular ssDNA in the isolate (Fig 5B).

## Discussion

While chemical and enzymatic in vitro synthesis methods remain popular for generating single-stranded DNA, leveraging filamentous phage biology offers a uniquely scalable alternative that shifts the primary bottlenecks from financial and enzymatic constraints to biological ones. By exploiting natural biological amplification within bacterial hosts, this system bypasses the prohibitive costs and kinetic thresholds of polymerases and nucleases typical of purely enzymatic approaches. The biological amplification inherent to phage production provides a practical route to scaling-up production, with further increases in culture volume potentially enabling higher total yields. With other cssDNA preparation methods this would involve increasing expensive enzyme inputs, dramatically lowering the technical and financial barriers when working with large constructs [8,26,27]. This scale capacity is particularly evident when handling multi-kilobase inserts, which frequently struggle to maintain structural integrity on synthetic platforms. The phage packaging mechanism readily accommodates these long sequences, rendering it highly effective for producing the large donor templates required in HDR and full-length gene replacement strategies [19,35,36]. Also, because the ssDNA is biologically encapsulated within a protein capsid during assembly and secretion, it gains intrinsic protection against mechanical shearing and enzymatic degradation [2,8]. This allows for the use of aggressive extracellular DNase treatments to eliminate contaminating host genomic and plasmid DNA before precipitation. When paired with anion-exchange chromatography, the workflow relies on charge-based separation to isolate the target nucleic acids, bypassing the harsh proteolytic treatments common to silica-based purification methods [37]. Because the entire process uses standard laboratory infrastructure and common Escherichia coli strains, it is easily adopted by standard molecular biology labs without the need for custom polymers or complex nick-and-digest systems [29,33,34].

cssDNA may be suitable for genome-editing applications as circular ssDNA donors have previously demonstrated advantages over linear donor formats by avoiding the activation of intracellular DNA damage responses and exhibiting superior stability [19,35].

Despite these clear operational benefits, the practical application of this protocol introduces distinct biological trade-offs, specifically the significant metabolic burden and sequence constraints imposed on the host by the requirement for phage-derived replication and packaging machinery, as well as sequence dependencies and processing constraints that warrant closer examination. The most prominent bottleneck is the protocol’s total reliance on filamentous phage biology, which restricts host selection to F-pilus-positive E. coli strains and requires compatible phagemid/helper systems. Specific target sequences, particularly those containing secondary structures, inverted repeats, or toxic domains, can actively disrupt viral packaging and host secretion kinetics, severely affecting yield or causing genetic drift during amplification [8,26,27]. To address this gap, future system improvements could explore the engineering of host strains with optimised chaperone expression to alleviate secretion stress or the utilisation of helper phages modified to tolerate structurally complex inserts.

Additionally, relying on biological fermentation extends the experimental timeline to 2–3 days, making the protocol inherently limited to single-batch processing compared to automated in vitro enzymatic or chemical methods that yield ssDNA within hours, albeit at a lower capacity and higher cost [28,34]. For projects where rapid turnaround is the priority rather than scale, this timeline may prove restrictive. While adapting the protocol to high-density bioreactor systems or continuous-flow fermentation models has been proposed to maximize volumetric productivity, these engineering approaches primarily improve overall biomass and yield rather than shortening the absolute biological timeline required for phage extrusion.

Beyond workflow kinetics, ensuring the structural integrity and purity of the final product demands more rigorous quality metrics than standard protocols typically require. Because standard phage replication does not inherently enforce rigid genetic boundaries on the final ssDNA length, a minor degree of length heterogeneity can occur [34,38]. While this structural variation rarely impairs basic HDR efficiency, users demanding absolute, single-nucleotide precision for sensitive downstream applications cannot rely solely on basic visual inspections via denaturing gel electrophoresis. Instead, incorporating automated electrophoresis platforms, such as a DNA TapeStation, is highly recommended to precisely assess size distribution, detect truncated fragments, and evaluate batch-to-batch structural integrity with high sensitivity. Furthermore, verifying exact sequence fidelity across the entire synthesized construct is paramount. Comprehensive Sanger sequencing using a series of overlapping walk-in primers should be integrated as a standard quality check to confirm that no mutations or deletions were introduced during biological replication. Similarly, confirming circular topology and the complete absence of host genomic or helper plasmid contamination is critical. While exonuclease I resistance assays provide a basic baseline [19,28], integrating quantitative PCR (qPCR) assays targeted at specific host and helper phage sequences would establish a far more robust threshold for trace double-stranded DNA contamination.

Finally, owing to the limitations mentioned above, there are several developments that could improve the performance and broader utility of this platform. Engineering the f1 initiation and termination elements is one such avenue, as modification of the pScaf terminator reduced, but did not completely eliminate, alternative start–stop products [37]. Larger mutational screens could therefore be employed to identify variants of these elements that retain efficient replication while producing a more homogeneous cssDNA product.

Production hosts could also be refined. Arabinose-inducible helper-phage-free systems and strains carrying chromosomally integrated M13 functions have shown that cssDNA production can proceed without conventional helper-phage infection [31,32]. The eScaf study identified construct-dependent differences in yield, purity and compatibility between phagemids and helper strains, supporting the need for development of sequence-aware design rules for optimal cssDNA production. These rules would take into account construct length, GC content, repeated regions, secondary structure and host toxicity. Generating larger datasets linking these features to production outcomes could support predictive models in identifying difficult constructs before cloning. [32].

For HDR applications, manufacturing attributes including the fraction of the full-length product, topology, residual dsDNA and endotoxin presence may influence knock-in efficiency, cell viability and inflammatory signalling. Published studies show that cssDNA performance varies across donor designs, nucleases, loci and cell types [9,19,20]. Letort et al. also observed inflammatory pathway activation in cssDNA-edited HSPCs and proposed testing inhibitory chemical modifications or gapmer sequences for future investigation to reduce innate immune sensing [20,39].

## Methods

### Materials/Reagents

#### Bacterial Culture

- E. coli XL1-Blue competent cells **(Generated in this lab)**
- M13 Helper Plasmid **(Addgene, Cat. 120340)**
- Phagemid vector containing desired insert and M13 origin **(Antibody Design Laboratories, PD0111)**
- 2×YT broth (16 g tryptone, 10 g yeast extract, 5 g NaCl per liter) **(HIMEDIA, M1251)**
- Appropriate antibiotic solutions i.e Carbenicillin, Kanamycin **(SRL Sisco Research Laboratories Pvt. Ltd.)**
- LB Agar **(HiMedia, G1151)**
- LB Broth **(HiMedia, M1245)**

#### Phage Harvesting

- DNase I (RNase-free) **(NEB, M0303S)**
- 1M MgCl₂ **(Sigma, Cat. M8266)**
- Ethylenediaminetetraacetic acid (EDTA), 0.5 M, pH 8.0 **(HiMedia, MB011)**
- Polyethylene glycol 8000 (PEG-8000) **(SRL Sisco Research Laboratories Pvt. Ltd., Cat. 44766)**
- Sodium chloride (NaCl) **(MP biomedicals, 194848)**

#### Phage Lysis

- Triton X-100 **(MP biomedicals, 194854)**
- Guanidine-HCl **(MP biomedicals, 194826)**
- MOPS (MP biomedicals, 102370)
- SDS (10% stock) **(USBiological Lifesciences, 151-21-3)**

#### cssDNA Isolation

- Proteinase K (20 µg/µL stock) **(Thermo Fisher, 100005393)**
- Phenol:Chloroform:Isoamyl Alcohol (25:24:1, PCI) **(Thermo Fisher, 15593031)**
- Isopropanol **(Merck supelco, 109634)**
- Chloroform (molecular biology grade) **(Merck, C2432)**
- 3 M Sodium acetate (pH 5.2–5.5) **(Merck, S2889)**
- Absolute ethanol (100%, chilled) **(Merck supelco, 1.00983)**
- 70% ethanol **(Merck supelco, 1.00983)**
- Qiagen QIAprep MaxiPrep Kit with QBT, QC, and QF buffers (or equivalent column-based purification kit) **(Qiagen, 12163)**
- Nuclease-free water (NFW) **(Ambion, AM9937)**

#### QC Tests

- Molecular Grade Agarose **(HiMedia, 9012-36-6)**
- ExoI nuclease **(NEB, M0568)**
- DpnI **(NEB, R0176S)**

### Equipment

- Shaker incubator capable of maintaining 25–37 °C
- UV Spectrophotometer
- Fixed Angle Centrifuge (4,000–15,000 × g)
- Swing-bucket rotor for large-volume centrifugation (4,000 x g)
- 2 L baffled Erlenmeyer flasks
- 0.22 µm syringe filter
- Water bath or heating block capable of maintaining 37 °C, 42 °C, 75 °C, and 80°C
- Agarose gel electrophoresis apparatus
- Microcentrifuge tubes (MCTs)
- Pipettes and sterile tips
- Vortex mixer (optional)
- QIAGEN-tip 500 column (or equivalent large-capacity anion-exchange column) **(Qiagen, 12163)**

### Buffers

- Buffer M1 and M2 i.e. precipitation and Phage lysis buffer to be prepared as follows:

- Buffer M1: Polyethylene glycol (PEG 8000); 3 M NaCl (Storage at 4°C)
- Buffer M2: 1% Triton X-100; 500 mM Guanidine-HCl; 10 mM MOPS, pH 6.5 (Storage at room temperature)
- TE Buffer (pH 8.0)

## Protocol

### 1. Transformation and Starter Culture Preparation

#### 1.1 Preparation of competent cells

*E. coli* XL1-Blue cells were prepared using the Inoue method (other F-pilus–positive strains with transformation efficiency ≥7 × 10⁷ cfu/µg DNA may also be used).

#### 1.2 Co-transformation

Competent cells were co-transformed with the M13 helper plasmid and the phagemid carrying the desired insert. The transformed cells were plated on selective agar containing the appropriate antibiotics to maintain both constructs and incubated at 30 °C for 16-19 h.

#### 1.3 Colony selection and clonal expansion

A single colony was picked and re-streaked (e.g., by quadrant streaking) to ensure clonal uniformity. The plate was incubated at 30 °C (or 25–30 °C) overnight, or until well-isolated colonies were obtained.

### 2. Large-Scale Culture and Phage Amplification

#### 2.1 Media composition

Autoclaved 2×YT medium containing 1.6% (w/v) tryptone, 1.0% (w/v) yeast extract, and 0.25% (w/v) NaCl was used.

#### 2.2 Culture scale-up

Large-scale culture was initiated by inoculating a single colony into 500mL-1L of 2×YT medium in baffled flasks and supplemented with the appropriate antibiotics.

#### 2.3 Growth conditions

Cultures were incubated with shaking at 180 rpm and 30 °C for 16 h, after which the temperature was increased to 37 °C and the shaking speed to 220 rpm. Although amplification can be performed entirely at 37 °C, incubation at a reduced temperature during the initial stages was used to limit rapid bacterial overgrowth and premature host-cell lysis, which could adversely affect phage recovery and cssDNA yield.

#### 2.4 Optimal harvest point

Cultures were harvested at an OD₆₀₀ of approximately 0.9–1.0. This optical density was selected to maximise phage production while minimising cell lysis and the release of contaminating nucleic acids [28].

### 3. Harvesting and Clarification of Phage-Containing Supernatant

#### 3.1 Cell removal by centrifugation

Cultures were centrifuged at 8,000–10,000 × *g* for 15–20 min at room temperature to pellet the bacterial cells.

***Optional:*** *In-process plasmid miniprep to check for presence of cssDNA*

#### 3.2 Supernatant transfer and clarification

The phage-containing supernatant was carefully transferred to fresh tubes without disturbing the bacterial pellet. The supernatant was centrifuged again, if necessary, to remove any residual cellular debris.

#### 3.3 Filtration

The clarified supernatant was filtered through a 0.22 µm membrane to remove any remaining bacterial cells.

#### 3.4 Pause point

The filtered supernatant could be stored at 4 °C overnight without any detectable loss of cssDNA yield or quality.

### 4. Nuclease Treatment

The ssDNA encapsulated within M13 phage particles is protected from DNase I digestion, whereas free ssDNA and dsDNA released from lysed *E. coli* cells or residual plasmid DNA are degraded.

#### 4.1 DNase I digestion

DNase I was added to a final concentration of 1 U/mL and MgCl₂ was added to a final concentration of 10 mM.

e.g. For 100 mL of culture supernatant:

- DNase I (2 U/µL): 50 µL
- 1 M MgCl₂: 1 mL

The samples were incubated in a water bath at 37 °C for 1 h with gentle inversion every 10 min.

#### 4.2 DNase inactivation

To terminate the reaction, EDTA was added to a final concentration of 10 mM.

e.g. For 100 mL of culture supernatant:

- 0.5 M EDTA (pH 8.0): 2 mL

### 5. Phage Precipitation and Lysis

#### 5.1 PEG/NaCl precipitation

Chilled PEG/NaCl precipitation buffer (Buffer M1; 0.2 volumes) was added to the supernatant, mixed thoroughly, and incubated on ice (4 °C) for 60 min.

#### 5.2 Phage pelleting

The precipitated phage particles were pelleted by centrifugation at 10,000 × *g* for 10 min at room temperature. A white translucent pellet, indicative of successful phage precipitation, was observed (Fig. 3B). The supernatant was carefully discarded, and the tubes were briefly centrifuged again (∼30 s) to collect and remove any residual liquid.

#### 5.3 Phage lysis

The phage pellet was resuspended in Buffer M2 and incubated at 80 °C for 40 min to lyse the phage particles and release cssDNA. The lysate was allowed to cool to room temperature and was mixed gently if phase separation was observed.

### 6. cssDNA Purification by Anion-Exchange Chromatography

#### 6.1 Column equilibration

The QIAGEN-tip 500 column was equilibrated with 10 mL of Buffer QBT.

#### 6.2 Sample loading

The lysed phage sample was loaded onto the QIAGEN-tip 500 column by gravity flow. If the lysate appeared cloudy, it was clarified by centrifugation or filtration prior to loading.

#### 6.3 Washing and elution

The column was washed with 2 × 30 mL of Buffer QC, and cssDNA was eluted with 15 mL of Buffer QF.

#### 6.4 DNA precipitation and resuspension

The eluted DNA was precipitated by adding 0.7 volumes of isopropanol and centrifuged for 30 min at ≥10,000 × *g*. The DNA pellet was washed with 70% ethanol, briefly air-dried, and resuspended in nuclease-free water or a Tris-based buffer.

#### 6.5 Quantification and quality assessment

The concentration and purity of the purified cssDNA were determined by UV spectrophotometry. The integrity of the isolated cssDNA was assessed by agarose gel electrophoresis using an appropriate gel concentration.

The identity and sequence fidelity of the produced cssDNA can further be validated by Sanger sequencing using primers designed across the cssDNA sequence.

#### 6.6 Quality Control

The circular topology of cssDNA can be assessed based on its resistance to exonuclease-mediated degradation. The purified cssDNA was evaluated for resistance to end-directed exonucleases, such as Exonuclease I, which preferentially degrade linear single-stranded DNA (Fig 6B). Additional treatment with single-strand-specific endonucleases, such as S1 nuclease or mung bean nuclease. These analyses should be performed alongside appropriate linear ssDNA and double-stranded plasmid DNA controls to enable accurate interpretation of nuclease sensitivity patterns.

**Figure 6.**
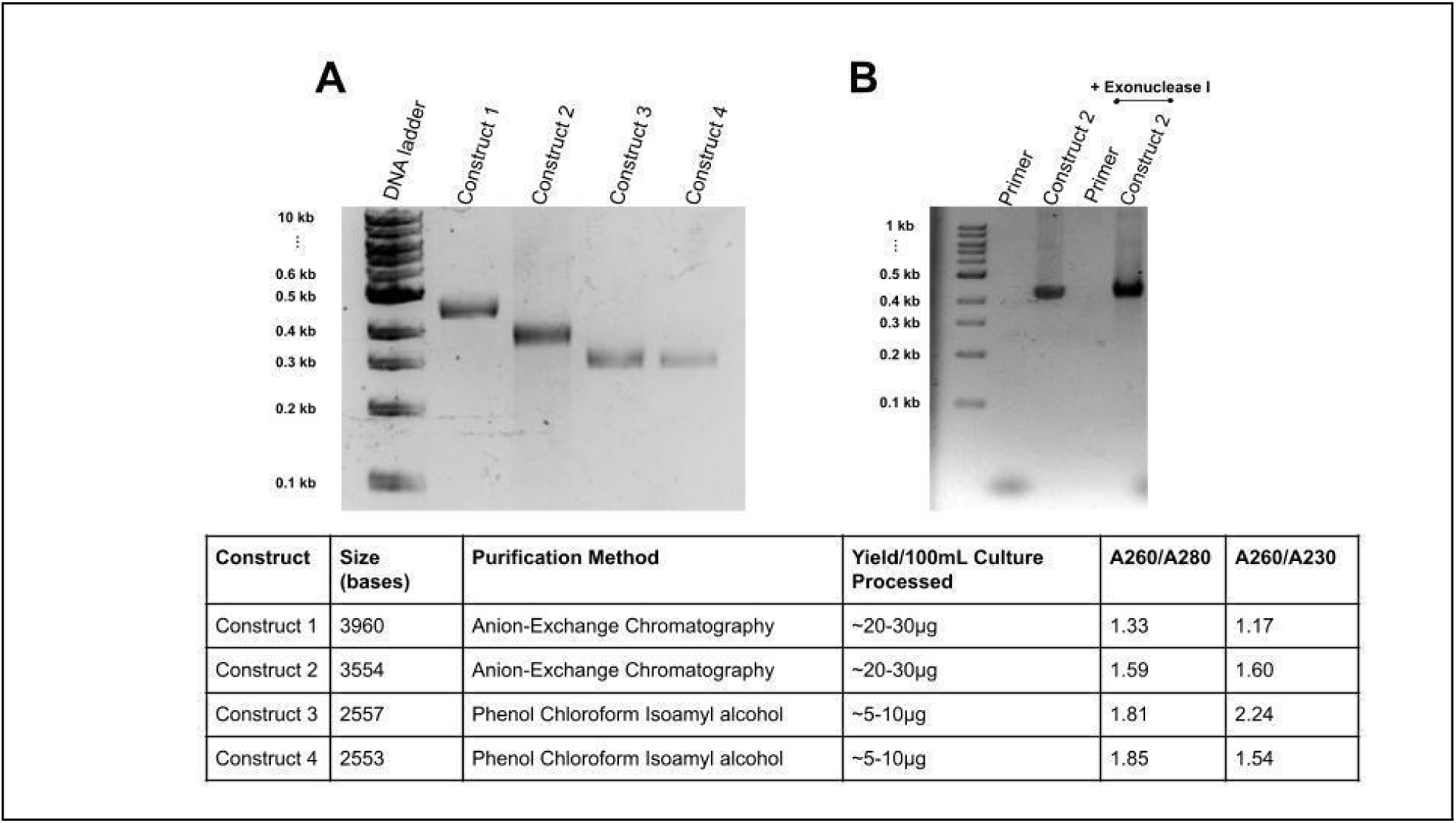
Characterisation of purified cssDNA isolates. **(A)** Representative agarose gel electrophoresis (1% agarose) of four different cssDNA constructs purified using two independent isolation methods. The corresponding yield and purity metrics for each isolate are summarised in the table below **(B)** Exonuclease I digestion was performed to assess the presence of contaminating linear single-stranded DNA (ssDNA). PCR primers, which serve as linear ssDNA substrates, were included as a positive control to confirm Exonuclease I activity.

### 7. Alternative to Anion-Exchange Chromatography - Purification by Phenol Chloroform Isoamyl alcohol

#### 7.1 Proteinase K Treatment

The PEG-precipitated phage pellet was resuspended in TE buffer (pH 8.0). SDS was added to a final concentration of 0.5%, followed by Proteinase K (50–100 µg/mL). The sample was incubated at 37 °C for 30–60 min to digest phage coat proteins and release cssDNA.

#### 7.2 PCI Extraction (25:24:1)

An equal volume of phenol:chloroform:isoamyl alcohol (PCI; 25:24:1) was added to the lysate. The sample was mixed gently by inversion and centrifuged at 12,000 rpm for 10 min at room temperature. The upper aqueous phase was carefully transferred to a fresh tube. The PCI extraction was repeated once, followed by a final chloroform extraction to remove residual phenol.

#### 7.3 Ethanol Precipitation

Sodium acetate (0.1 volume) and 2–2.5 volumes of chilled 100% ethanol were added to the aqueous phase. The sample was mixed gently and incubated at −20 °C for 1–2 h or overnight to precipitate DNA.

#### 7.4 DNA Pelleting

The precipitated DNA was collected by centrifugation at 12,000–14,000 rpm for 15–20 min at 4 °C. The supernatant was carefully removed without disturbing the DNA pellet.

#### 7.5 Ethanol Wash

The DNA pellet was washed with 0.5–1 mL of 70% ethanol and centrifuged at 6,000 rpm for 10 min. The supernatant was carefully discarded, and residual ethanol was removed.

#### 7.6 Drying and Resuspension

The DNA pellet was briefly air-dried for 5–10 min to remove residual ethanol, avoiding complete drying of the pellet. The purified DNA was resuspended in approximately 50 µL of nuclease-free water or TE buffer.

### Troubleshooting guide

**Table 2:**
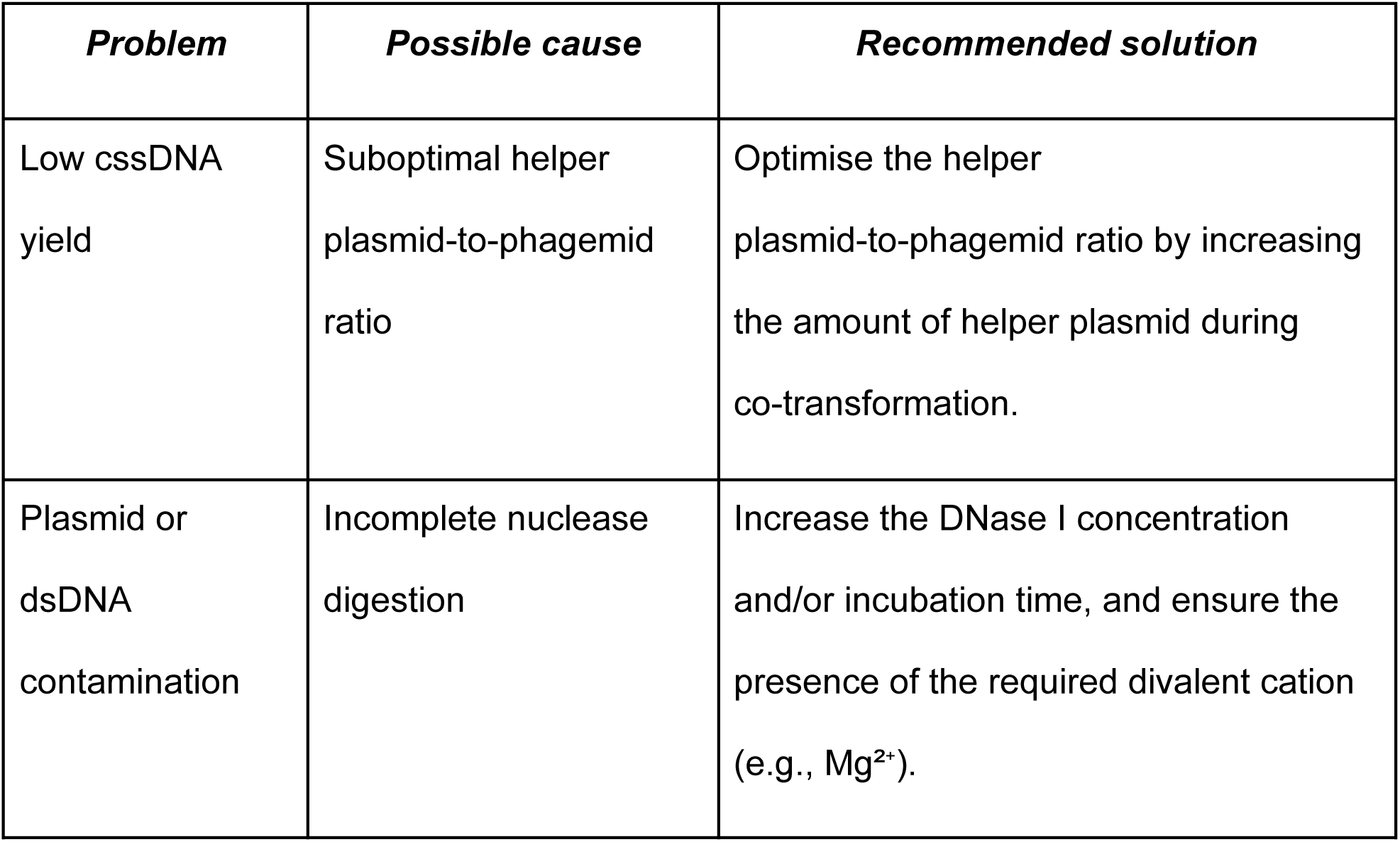
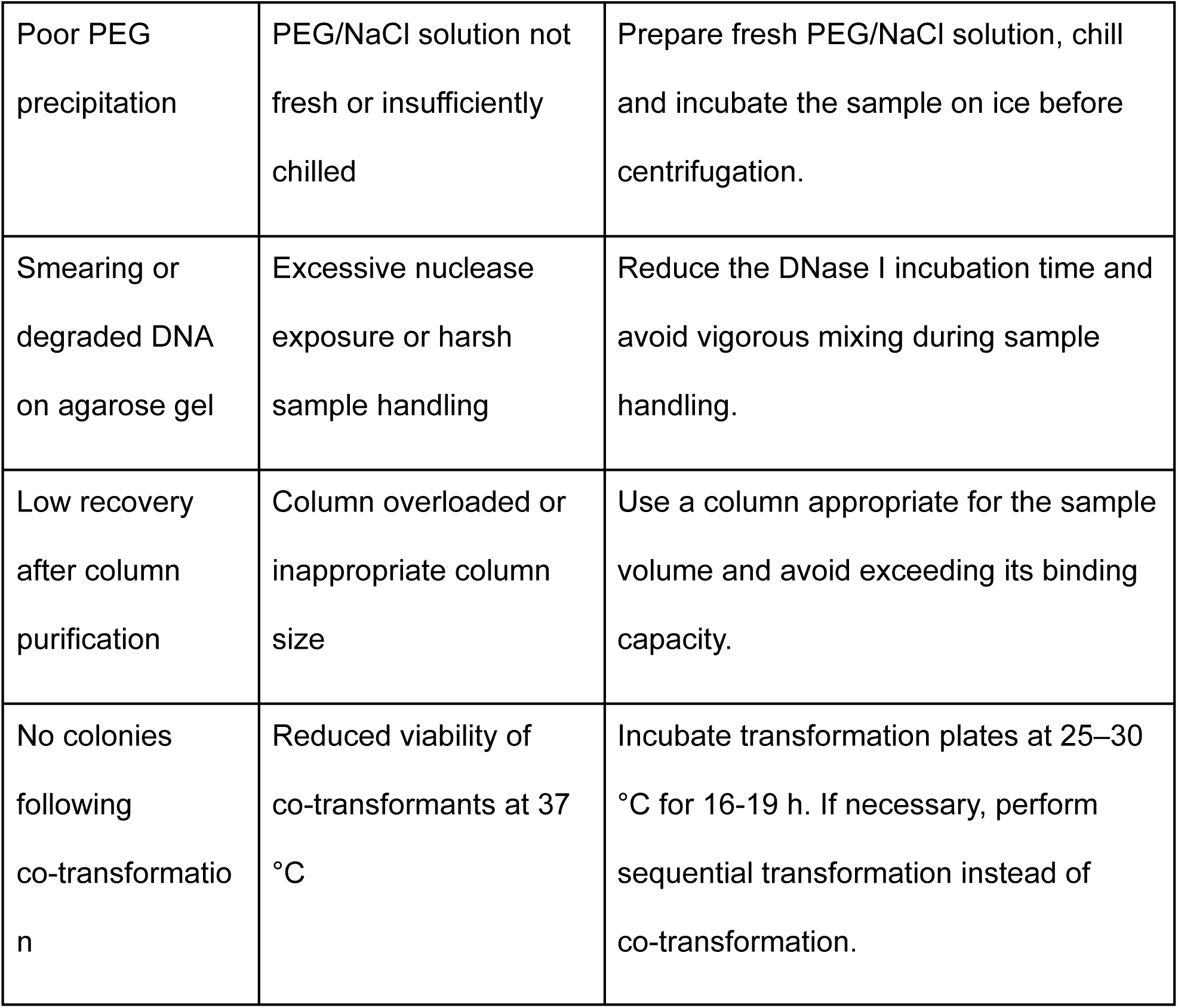
Common issues encountered during cssDNA production and suggested remedies.

## Conclusion

Collectively, this protocol provides a robust, reproducible, and scalable approach for preparative-scale cssDNA production using standard molecular biology infrastructure, with the potential to support research and translational downstream applications.

## Author contribution

**Susi Mathews:** Conceptualization, Methodology, Investigation, Project administration, Writing - Original Draft, Writing – Review & Editing

**Mahima Kapoor:** Conceptualization, Methodology, Investigation, Writing - Original Draft

**Akschya Sivacoumar:** Methodology, Investigation, Writing - Original Draft

**Raj Acharya:** Methodology, Investigation, Validation, Writing - Original Draft

**Debojyoti Chakraborty:** Conceptualization, Resources, Funding acquisition, Supervision

## Declaration of Generative AI and AI-Assisted Technologies in the Manuscript Preparation Process

During the preparation of this work, the authors used ChatGPT to improve the language and readability of the manuscript. The authors reviewed and edited the output as needed and take full responsibility for the content of the published article.

## Acknowledgement

This study was funded by the grant, ‘CRISPR-Cas9 Technology Implementation and Support Activities to Start-ups and MSME’ (OLP242503 (CSIR-IGIB)), and the infrastructure for the work was provided by CSIR-Institute of Genomics and Integrative Biology (CSIR-IGIB).

